# Chaotic provinces in the kingdom of the Red Queen

**DOI:** 10.1101/062349

**Authors:** Hanna Schenk, Arne Traulsen, Chaitanya S. Gokhale

## Abstract

The interplay between parasites and their hosts is found in all kinds of species and plays an important role in understanding the principles of evolution and coevolution. Usually, the different genotypes of hosts and parasites oscillate in their abundances. The well-established theory of oscillatory Red Queen dynamics proposes an ongoing change in frequencies of the different types within each species. So far, it is unclear in which way Red Queen dynamics persists with more than two types of hosts and parasites. In our analysis, an arbitrary number of types within two species are examined in a deterministic framework with constant or changing population size. This general framework allows for analytical solutions for internal fixed points and their stability. For more than two species, apparently chaotic dynamics has been reported. Here we show that even for two species, once more than two types are considered per species, irregular dynamics in their frequencies can be observed in the long run. The nature of the dynamics depends strongly on the initial configuration of the system; the usual regular Red Queen oscillations are only observed in some parts of the parameter region.

## 1 Introduction

Studying host-parasite coevolution using mathematical models has lead to substantial advances in our understanding of the dynamics of the interaction. For example hypothesising the role of reciprocal selection between the antagonistic species in the evolution of virulence and tolerance. We specifically focus on the Red Queen hypothesis (van Valen, 1973; Stenseth and Maynard Smith, 1984; Dieckmann et al., 1995; Clay and Kover, 1996; Salathé et al., 2008). The hypothesis has been used in a broad context, leading to multiple definitions (Brockhurst et al., 2014; Rabajante et al., 2015). According to Van Valen, the maintenance of biodiversity is possible as long as the species displace each other, or when the resource distribution changes over time (van Valen, 1973). However, the different definitions are underlined by the presence of the typical dynamics expected within a species, namely Red Queen oscillations. These oscillations imply an interaction where the increase in the relative abundance of a a certain type within a species indicates an equal decrease in relative abundance of another type (Maynard Smith, 1976; Van Valen, 1977). In the context of hosts and parasites, indications for such oscillations in densities have been empirically confirmed for e.g. in dormant stages of the water flea *Daphnia magna* from pond sediments (Decaestecker et al., 2007) and freshwater snails *Potamopyrgus antipodarum* (Koskella and Lively, 2009). However, while it is already difficult to analyse these dynamics experimentally over a single cycle, the long term dynamics of such systems is challenging.

The co-evolution of hosts and parasites has for example been used to explain sexual reproduction (Lively, 2010). However, when multilocus genetics is at play, features of co-evolution models, such as the maintenance of polymorphism and evolution of sex, depend on the exact interactions patterns (Frank, 1993a; Parker, 1996; Frank, 1996; Sasaki, 2000; Metzger et al., 2016). Exploring a variety of interaction patterns between multiple types of hosts and parasites, we show that shortterm oscillations as the ones observed experimentally can be recovered in virtually all of these models, but one has to be very careful in extrapolating this kind of dynamics over a wider time horizon.

Mathematical models of host-parasite interactions with two types have been extensively analysed. A specific experiment in *Daphnia magna* (Carius et al., 2001) showed considerable variation in hosts (susceptibility) and parasites (infectiousness) of nine distinct types, which illustrates the necessity of considering models with more than two types. A recent model (Rabajante et al., 2015) based on ordinary differential equations numerically explored different numbers of types. The result strengthened the theory of the oscillatory Red Queen dynamics. In such numerical models, a broader exploration of the parameter space can lead to more general results and show the robustness of models. Here, we take a different approach and ask how complicated a model can become before the regular frequency dependent oscillations are lost? To tackle this question, we used analytical tools in addition to numerical integration.

Another aspect to be considered is the impact of population size (Papkou et al., 2016): A host population suffering from intense parasite pressure should decrease in absolute size. Similarly, a parasite population not finding sufficient hosts should decrease in absolute size. Recently, the impact of such changing population sizes was studied for two types (Gokhale et al., 2013; Song et al., 2015), including matching alleles (Frank, 1993b) or the gene-for-gene type of interactions (Flor, 1955; Engelstädter, 2015; Agrawal and Lively, 2002). By adjusting the birth rates of hosts and death rates of parasites to include frequency dependence, we can impose a constant population size. Such a transformation makes the underlying model identical to replicator dynamics (Taylor and Jonker, 1978; Hofbauer and Sigmund, 1998; Schuster and Sigmund, 1983).

Here, we extend the two approaches with changing and constant population size to an arbitrary number of types of hosts and parasites. As a simple example, we first consider the matching allele model: Each host can only be infected by its specific parasite type, and each parasite can only affect the one matching host. Next, cross-infectivity is incorporated so that two genetically similar parasites (neighbours to the focal parasite) can additionally infect a particular host in an equally robust manner and vice versa: each parasite can infect not only one host but also two more closely related hosts. Finally, a general model where different infectivity magnitudes are realised for each parasite type is analysed.

## 2 Model

### 2.1 Interactions between hosts and parasites

The number of parasites affecting a focal host (and vice versa) and the strength of the interactions are key components for models of host-parasite coevolution. Three possible models are depicted in Table 1, where fitness effects are collected in a matrix, which intuitively describes the influence of each type within one species on each type within the other species. Assuming *n* types of hosts and *n* types of parasites, *M^H^* describes the average loss of fitness hosts suffer from specific parasite types and *M^P^* describes the average gain of fitness parasites extract from the interaction. For example (*M^H^*)_2,4_ is the fitness loss that host 2 suffers from parasite type 4. On the other hand, (*M^P^*)_4,2_ is what parasite 4 gains from host 2.

To introduce host-parasite dynamics, we focus on the matching allele model first (Grosberg and Hart, 2000; Carius et al., 2001), where only fixed pairs of hosts and parasites can directly interact with each other (Tab. 1, Matching alleles). Interactions with all other partners are neutral and do not influence fitness. In a cross-infection model, it is instead assumed that neighbouring parasite types are genotypically or phenotypically similar in their infectiveness (Tab. 1, Cross-infection). This also applies to each host and its neighbours which have not developed resistance and are therefore susceptible to a specific parasite type which now benefits from three host types. In our model, we assume that cross-infectivity follows periodic boundary conditions where types 1 and *n* can also interact with three types of the other species. Finally, in our most general model hosts have a positive effect *α_i_* on parasites which have a negative effect –*cα_i_* with *c* > 0 on the hosts (Tab. 1, General infection). Every diagonal has a different value, which leads to *n* interaction parameters. This means that parasite *i* has the same effect –*cα_i_* on host *i* + *k* as parasite *j* on host *j* + *k*. Further, host *i* has the same but positive effect *α_i-k_* on parasite *i* – *k*. The restriction *M^H^* = –*c* ·(*M^P^*)^T^ ensures a scaled effect of the interaction partners. In this way we can for example envision a scenario in which there are matching hosts and parasites (the main diagonal) and the effect they exert on each other declines with distance between them, *α*_1_ > *α*_2_ > *α*_3_ >….

**Table 1:** Interaction models: *M^P^* is the parasite’s (row) gain achieved by a specific host (column). *M^H^* is the host’s (row) loss by a parasite (column).

| Model | $M^P$ | $M^H$ |
| --- | --- | --- |
| Matching alleles | $\begin{pmatrix} +1 & 0 & \cdots & 0 \\ 0 & +1 & \cdots & 0 \\ \vdots & \vdots & \ddots & \vdots \\ 0 & 0 & \cdots & +1 \end{pmatrix}$ | $\begin{pmatrix} -1 & 0 & \cdots & 0 \\ 0 & -1 & \cdots & 0 \\ \vdots & \vdots & \ddots & \vdots \\ 0 & 0 & \cdots & -1 \end{pmatrix}$ |
| Cross-infection | $\begin{pmatrix} +1 & +1 & 0 & 0 & \cdots & +1 \\ +1 & +1 & +1 & 0 & \cdots & 0 \\ 0 & +1 & +1 & +1 & \cdots & 0 \\ \vdots & \vdots & \vdots & \vdots & \ddots & \vdots \\ 0 & 0 & \cdots & +1 & +1 & +1 \\ +1 & 0 & \cdots & 0 & +1 & +1 \end{pmatrix}$ | $\begin{pmatrix} -1 & -1 & 0 & 0 & \cdots & -1 \\ -1 & -1 & -1 & 0 & \cdots & 0 \\ 0 & -1 & -1 & -1 & \cdots & 0 \\ \vdots & \vdots & \vdots & \vdots & \ddots & \vdots \\ 0 & 0 & \cdots & -1 & -1 & -1 \\ -1 & 0 & \cdots & 0 & -1 & -1 \end{pmatrix}$ |
| General infection | $\begin{pmatrix} \alpha_1 & \alpha_2 & \cdots & \alpha_{n-1} & \alpha_n \\ \alpha_n & \alpha_1 & \alpha_2 & \cdots & \alpha_{n-1} \\ \vdots & \alpha_n & \alpha_1 & \cdots & \vdots \\ \alpha_3 & \vdots & \vdots & \ddots & \alpha_2 \\ \alpha_2 & \alpha_3 & \cdots & \alpha_n & \alpha_1 \end{pmatrix}$ | $\begin{pmatrix} -c\alpha_1 & -c\alpha_n & \cdots & -c\alpha_3 & -c\alpha_2 \\ -c\alpha_2 & -c\alpha_1 & -c\alpha_n & \cdots & -c\alpha_3 \\ \vdots & -c\alpha_2 & -c\alpha_1 & \cdots & \vdots \\ -c\alpha_{n-1} & \vdots & \vdots & \ddots & -c\alpha_n \\ -c\alpha_n & -c\alpha_{n-1} & \cdots & -c\alpha_2 & -c\alpha_1 \end{pmatrix}$ |

We stress that these matrices are not chosen to represent a particular biological system. Instead, our approach is to consider more complex models beyond the gene for gene or matching allele models and to analyse their dynamics. The fitness effects represented in the matrices now have to be included in our models. We start with a changing population size approach and then turn to constant population size.

### 2.2 Changing population size

The classical Lotka–Volterra dynamics are usually employed to describe predator-prey systems where the prey reproduces at a constant rate and the predator dies at a constant rate (Lotka, 1925; Volterra, 1928). This allows the population size to change. The predator density is influenced by the abundance of prey and the prey density is influenced by the abundance of predators. The same concept can be applied to host-parasite systems. We assume *n* different types of hosts and *n* different types of parasites. The hosts have a constant birth-rate *b*_*h*_ and a death rate that is determined by the interactions with the parasite. Conversely, we assume a constant parasite death-rate *d*_*p*_, but a birth rate that depends on the interactions with the hosts. With these assumptions the change of host (*h*_*i*_) and parasite (*p_i_*) abundance in time can be formulated as

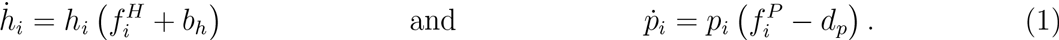

The fitness values 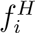 and 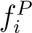 are defined by the interaction matrix and the abundances of the types.

Instead of immediately numerically exploring the dynamics for particular parameter sets, we first aim to obtain some general insight. On the boundaries of the state space, we have one fixed point where hosts and parasites are extinct, *h*_*i*_ = *p*_*i*_ = 0 for all *i*. In the absence of parasites, the host population will continue to increase in size, whereas a parasite population is not viable in the absence of hosts. In terms of co-existence, it is more interesting to consider potential interior fixed points.

For the matching allele model, we have a fixed point where all hosts and parasites have equal abundances, 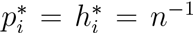 for all *i*. In this case, the equations completely decouple and each host-parasite pair can evolve independently. Thus, the fixed point is neutrally stable, as for the case of a single host and a single parasite.

For the cross-infection model, a host suffers from three parasite types and each parasite type benefits from three host types. The internal fixed point is now 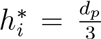 and 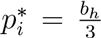. For the general model, we obtain 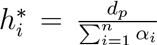 and 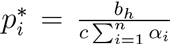. In both cases, we again find neutral stability (see supplementary material).

In addition, the symmetry of the system leads to a constant of motion. For all three models, the constant of motion is given by (Plank, 1995; Hofbauer and Sigmund, 1998),

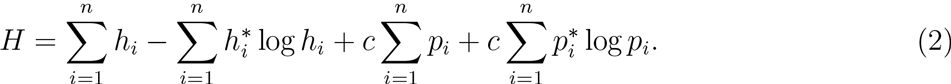

This implies that the dynamics effectively takes place in a space which has one dimension less.

### 2.3 Constant population size

As before we assume *n* different types of hosts and n different types of parasites. The relative abundance of host type *i* is *h_i_*, the relative abundance of parasite type *i* is *p_i_* (*i* = 1,…, n). With *h* and *p* we denote the vectors of the relative abundances. Thus, we have 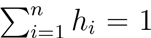 as well as 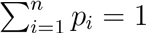. We assume that the relative abundances change according to the replicator dynamics (Hofbauer and Sigmund, 1998),

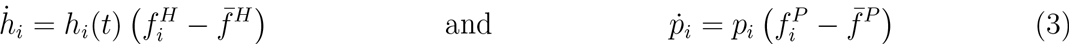

where 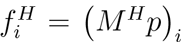 is the host fitness for type *i* and 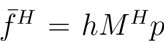 is the average fitness of the host population. Similarly, 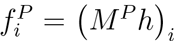 is the parasite fitness for type *i* and 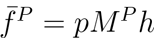 is the average fitness of the parasite population.

For example, a system with two hosts and two parasites (*n* = 2) where matching hosts and parasites have an influence of *α*_1_ = 1 and mismatching pairs exert a smaller fitness effect *α*_2_ = 0.3 with a twofold impact on the host *c* = 2 would be a system of four differential equations,

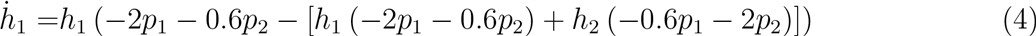

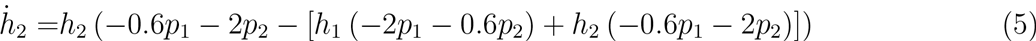

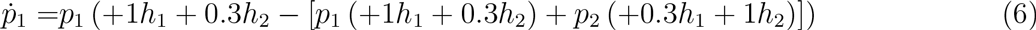

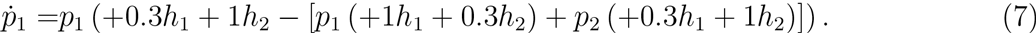

Again, this model can now be solved numerically to generate trajectories depending on the initial values of *p*_1_ and *h*_1_ (which determine *p*_2_ = 1 – *p*_1_ and *h*_2_ = 1 – *h*_1_). However, this approach would only lead to insights about particular parameter sets. Thus, here we take a different and–in our opinion–a more powerful approach and look at general properties of the system.

The replicator dynamics Eq. (3) has fixed points on the edge of the state space, e.g. *h*_1_ = *p*_1_ = 1 and *h_i_* = *p_i_* = 0 for *i* > 1. However, these fixed points are unstable for a generic parameter choice, which means that a small perturbation from this point drives the system away. There is an additional fixed point where all types have equal abundance, 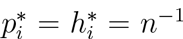 for all *i*. This arises from the symmetry of the interaction matrices we consider, but the fixed point can also be verified directly in Eq. (3).

The dynamics of the system depends crucially on the stability of the interior fixed point, which can be attracting, repelling or neutrally stable. For the matching allele model, the interior fixed point is neutrally stable for any number of types (see supplementary material), which implies that a small perturbation from the fixed point does not lead back to it, neither does it increase the distance. In terms of the cross infection models, it is substantially harder to prove this, but in the supplementary material we show that at least for *n* ≤ 6, the fixed point remains neutrally stable. Also for the general model, an analysis is intricate. For *n* = 3, we can show that the fixed point remains neutrally stable if the interaction strength decreases with the distance between host and parasite type.

There is a constant of motion, as recognized previously (Hofbauer, 1996)

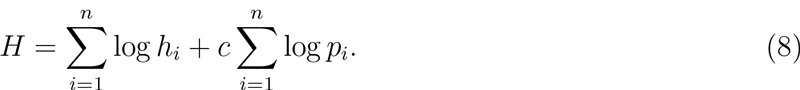

The existence of such a quantity arises from the symmetry of the system and implies that effectively, the system has one free variable less. However, as we show below, it does not imply any regularity of the dynamics.

### 2.4 Irregular dynamics in the most simple model

While the general properties discussed above lead to a first insight, e.g. the fact that there is always an interior fixed point and that it is neutrally stable, they do not give insights beyond the fact that the dynamics is oscillatory. Close to the fixed point, one would expect regular oscillations, but it remains unclear what happens if we leave the vicinity of the fixed point.

It turns out that in spite of the constants of motion and neutral stability, the trajectories of host and parasite abundances through time can become irregular and non-periodic. This can already be observed in a three type matching allele model with constant population size, cf. Fig. 1. This surprising result even for the simplest model we consider led us to examine this particular model more thoroughly.

**Figure 1:**
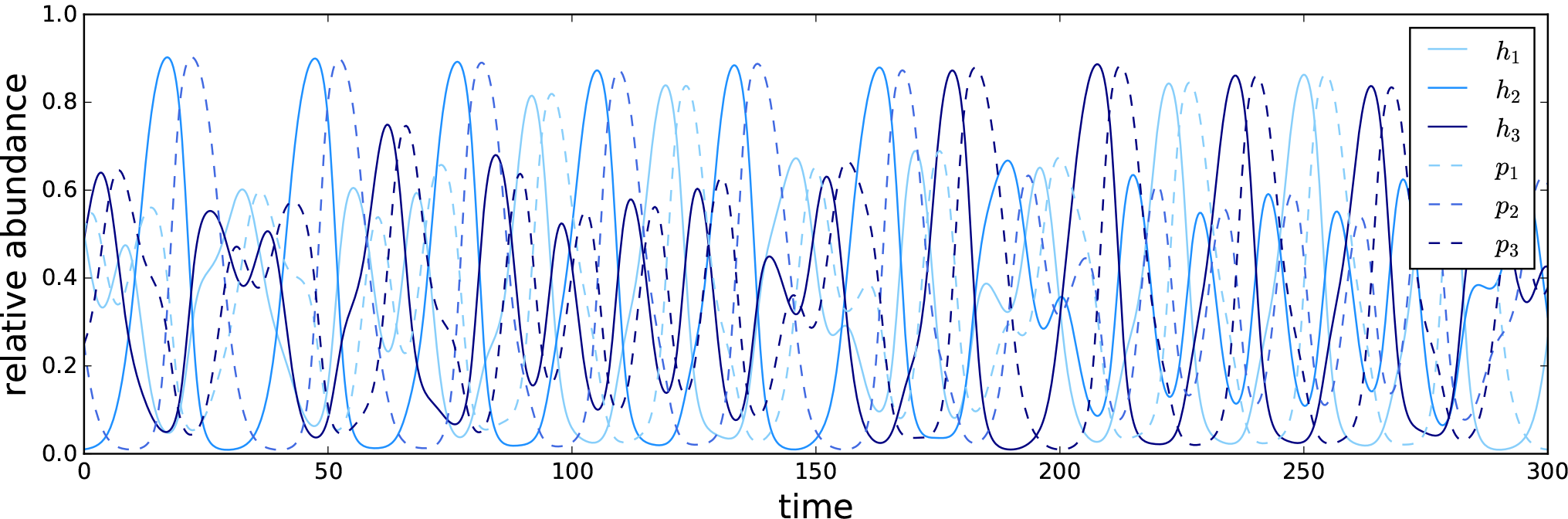
Matching allele replicator dynamics with three types: Trajectory of all host and parasite types for a 3-type matching allele replicator dynamics system, Eqs. (3), with initial conditions *h*(0) = (0.5, 0.01, 0.49)^T^ and *p*(0) = (0.5, 0.25, 0.25)^T^. Numerical integration with `python`’s built-in `odeint` function.

Because of the constant population size, a third type has a relative abundance determined by the abundance of the other two types. For each species the dimensions reduce from three to two. It is therefore possible to show the dynamics for one species in a 3-simplex (Figure 2), where each vertex represents the sole existence of one type, the edges correspond to a coexistence of two types and the interior is a state where no type is extinct. For balanced initial conditions close to the center of the simplex, trajectories are confined to orbits around the interior fixed point. For more extreme initial conditions, starting close to the edge of the simplex, this is no longer true. The trajectory is no longer limited to regular orbits, but nearly fills out the whole simplex, going from conditions close to extinction of one type (edges of simplex) to a near balance of all types (close to the interior fixed point).

To analyse the regularity of the dynamics further, we visualised trajectories of different initial conditions in Poincaré sections to check for chaotic behaviour (Strogatz, 2000). Plotting Poincare sections is a method to analyse dynamic properties of high dimensional systems. This is implemented by plotting trajectories in a two-dimensional area under certain restrictions (see Fig. 3). Periodic trajectories pass through the section in a periodic way, drawing circles or other closed trajectories. Chaotic trajectories have a much less ordered path and thus scatter over a larger part of the section. In Sato et al. (2002) chaotic behaviour and large positive Lyapunov exponents were found for several initial conditions in a two-person rock-paper-scissors learning game. This is formally closely related to a replicator dynamics host-parasite system with three types. We utilised this approach for our matching allele model and numerically evaluated several initial conditions. As expected from the neutrally stable fixed points, closed and periodic trajectories are found in Figure 3 for most initial conditions. For initial conditions close to the edge of the state space, the trajectories become visibly scattered.

**Figure 2:**
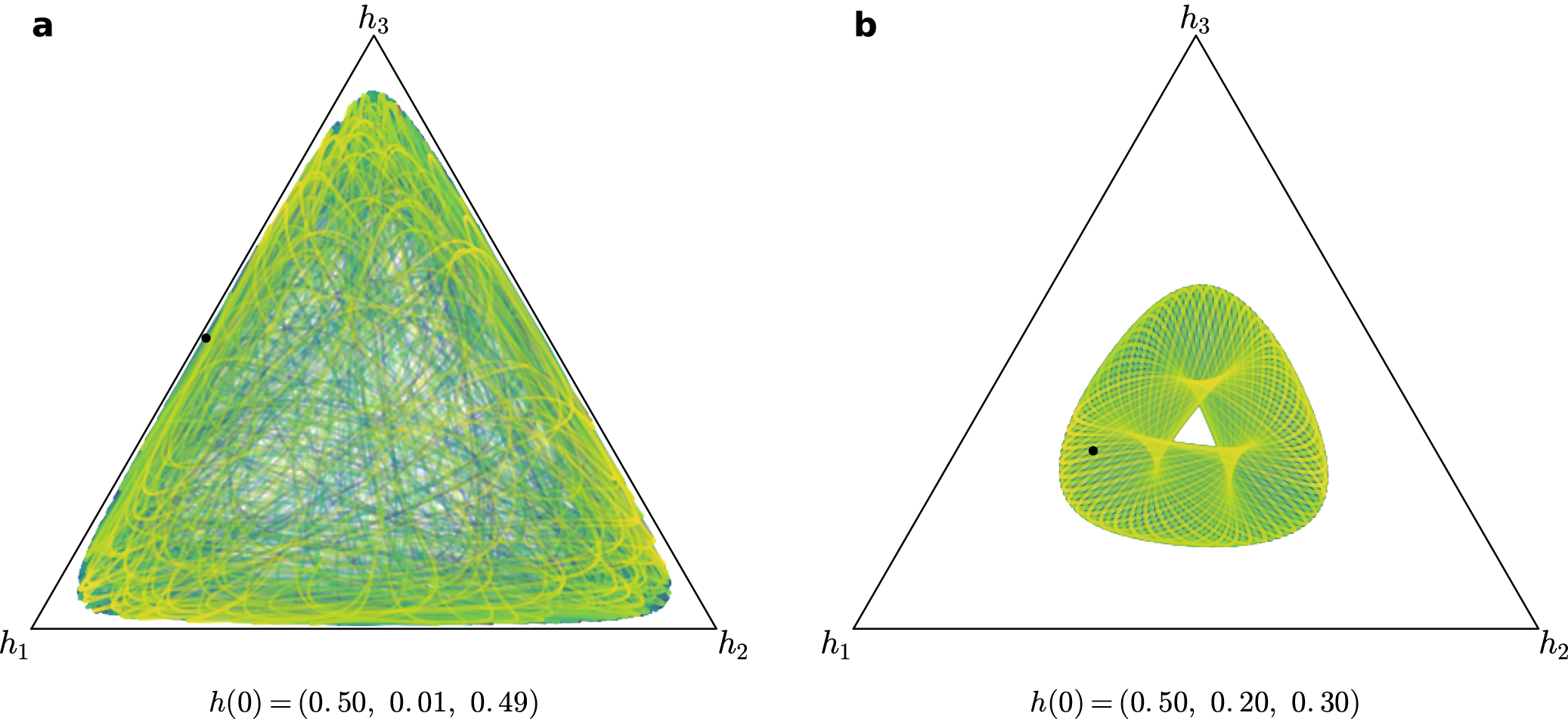
Matching allele replicator dynamics with three types. THost 3-simplex for a 3-type matching allele replicator dynamics system with initial conditions given by *h*(0) and *p*(0) = (0.5, 0.25, 0.25)^T^ indicated as a black dot. Panel (a) corresponds to the initial condition from Fig. 1. Time is represented in the colour gradient going from purple to blue, green and yellow. For initial conditions close to the interior fixed point, the dynamics remains regular (b), but for initial conditions closer to the edges, irregular dynamics emerges (a). Numerical integration was performed using `python`’s built in `odeint` function. Plotted for 10000 (a) and 5000 (b) generations.

**Figure 3:**
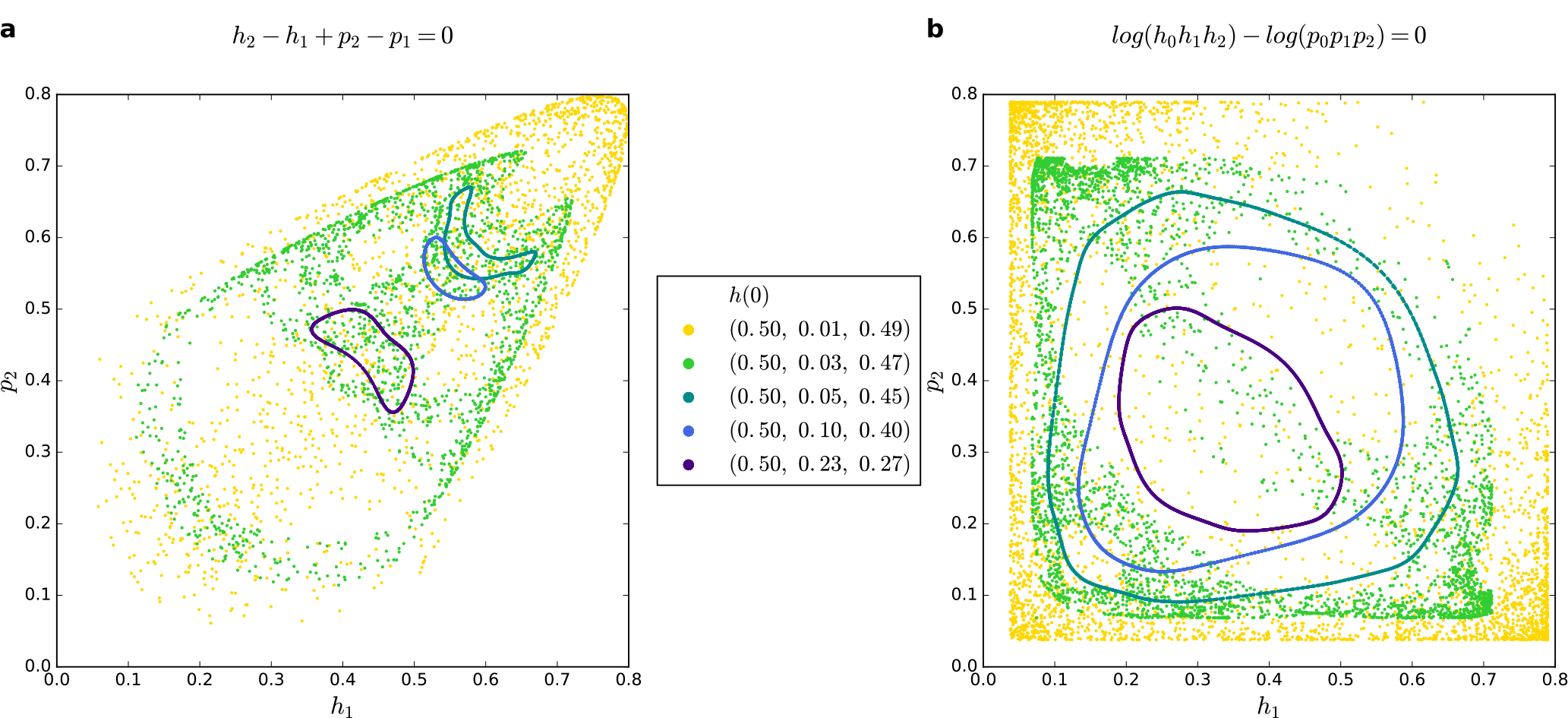
Poincaré sections for a 3-type matching allele replicator dynamics system: The Poincaré sections with restrictions |*h*_2_ – *h*_1_ + *p*_2_ – *p*_1_| ≤ 0.001 ((a), following Sato et al. (2002)) and | log *h*_1_*h*_2_*h*_3_ – log*p*_1_*p*_2_*p*_3_| ≤ 0.001 (b) are plotted. The horizontal and vertical axes are the host type 1, h1 and parasite type 2, *p*2 respectively. Initial conditions for *h*(0) are as stated in the legend and *p*(0) = (0.5, 0.25, 0.25)^T^. For initial conditions closer to the fixed point (dark green, blue, purple, see also Fig. 2b), the trajectories show periodic behaviour in a higher dimension. For extreme initial conditions, close to the edge of the state space (yellow, light green, see also Fig. 2a), the trajectories become chaotic and show a wide spread over the state space. Numerical integration was performed using `python`’s built in `odeint` function, for 50000 generations.

## 3 Discussion

Stenseth and Maynard Smith (1984) as well as Nordbotten and Stenseth (2016) showed that only trophic +/– interactions (as opposed to mutualism +/+ or competition –/–) promote Red Queen dynamics independent of abiotic factors. This justifies our study of these dynamics in a simple framework without abiotic influence or other types of interactions than trophic. Red Queen dynamics have been repeatedly reported to occur in models with two types, often because two alleles were in focus in the matching allele or gene-for-gene model (Schmid-Hempel and Jokela, 2002; Frank, 1993b; Flor, 1956; Agrawal and Lively, 2002; Song et al., 2015). A clear focus on multiple types has, to our knowledge, not been extensively analysed before. Yet, examples from observed biological systems clearly motivate the need for including this aspect into theoretical studies (Carius et al., 2001; Koskella and Lively, 2009; Luijckx et al., 2014). Rabajante et al. (2015) numerically investigated such host-parasite systems with multiple types. We study host parasite coevolution with three successively complicated interaction matrices in frameworks with more than two types where both constant and changing population size models can be justified (MacArthur, 1970).

We allow for *n* different interaction parameters in the most general payoff matrix. It is an advantage to be able to calculate specific outcomes with analytically derived statements and not have to rely entirely on numerical integration with fixed parameters and fixed initial conditions. For the same reason, we focus on a deterministic framework, allowing broad predictions. Eventually, one also has to include stochastic effects, which can have decisive impact on coevolutionary dynamics, in particular when the dynamics reaches the edges of the state space where extinction is likely (Gokhale et al., 2013). Also to explore signatures of genomic selection, such as selective sweeps or balancing selection, this is necessary (Tellier et al., 2014).

The presence of neutrally stable fixed points and constants of motion may lead to the belief that Red Queen dynamics exist on stable, regular orbits around the interior fixed point. These are also often shown to illustrate this kind of dynamics. The fixed points are neither repelling, thereby forbidding a coexistence of this type, nor attracting and, therefore, leading to a stable equilibrium. A neutrally stable fixed point and the consequent concentric circles, spheres or higher dimensional circulations around the point mean that the system is constantly changing, and yet, stationary in this change. Formulating constants of motions or Hamiltonians underlines this principle. However, the stability of a fixed point only holds locally and a constant of motion is a purely mathematical constraint. In general, the neutral stability supports the notion of Red Queen cycling. To understand this kind of dynamics in more detail, we have focused on the simplest possible dynamics and not attempted to construct a model for a concrete biological scenario. It is possible to consider more complex dynamics with stable or unstable interior fixed points or even limit cycles. However, our goal is to illustrate that even these simple models, which often form the basis for investigations of host parasite coevolution, can show a dynamics which is much richer than one would expect from verbal arguments or numerical considerations of such systems close to interior fixed points.

Simple models built on differential equations have been famously known to show chaotic properties in the sense that close by starting conditions can lead to very diverse outcome, thus restricting the predictability of the dynamics to very short time horizons (Lorenz, 1963; May, 1976; Hamilton et al., 1990; Hassell et al., 1991; Schuster, 1995; Sato et al., 2002). It thus comes as no surprise that a system of multiple interacting species can lead to chaos in some parts of the parameter space (May and Leonard, 1975; Smale, 1976). Recently Duarte et al. (2015) found chaos in a food chain model with three species resulting in Red Queen dynamics. There is general interest in increasing the number of species in the analysis (Liow et al., 2011; Dercole et al., 2010). However, to our knowledge, chaos has never been linked to Red Queen dynamics in a two-species model.

For multiple (three and more) types, we found that trajectories starting further away from the interior fixed point can show such chaotic behaviour. Chaotic fluctuations of host and parasite abundances, therefore, become possible in parts of the parameter space. A new type can be introduced to a system exhibiting typical Red Queen oscillations, e.g. via mutation or migration. While the mutant appears at low frequencies, the system shifts to an edge in a higher dimension. Our analysis predicts that this might often lead to chaotic dynamics rather than to stasis or the persistence of regular Red Queen oscillations. The typical, Red Queen dynamics is thought to consist of regular sinusoid-like oscillations of the frequencies of the different types within host and parasite populations with short periods and one or few amplitudes. We are now facing highly irregular trajectories without periodic re-occurrence and different magnitudes of maxima in each cycle. With our model, we propose that in addition to the concepts of stasis or regular Red Queen cycling a third scenario-chaotic Red Queen dynamics-is possible and likely. Chaos, then, would be especially rampant in the presence of low levels of standing genetic variation, mutations and migration. Moreover, it would in particular occur for very large populations, where the typical intuition of evolutionary biologists is to expect regular deterministic dynamics.

## Acknowledments

We thank Hinrich Schulenburg for fruitful discussions and comments on the manuscript. Generous funding by the Max Planck Society is gratefully acknowledged. CSG acknowledges funding from the New Zealand Institute for Advanced Study and support from the Marsden Fund Council administered by the Royal Society of New Zealand. The authors declare no conflict of interests.

